# Colorectal cancer progression to metastasis is associated with dynamic genome-wide biphasic 5-hydroxymethylcytosine accumulation

**DOI:** 10.1101/2025.02.21.639484

**Authors:** Ben Murcott, Floris Honig, Dominic Oliver Halliwell, Yuan Tian, James Lawrence Robson, Piotr Manasterski, Jennifer Pinnell, Thérèse Dix-Peek, Santiago Uribe-Lewis, Ashraf EK Ibrahim, Julia Sero, David Gurovich, Nikolas Nikolai, Adele Murrell

## Abstract

**Background:** Colorectal cancer (CRC) progression from adenoma to adenocarcinoma is associated with global reduction in 5-methylcytosine (5mC) and 5-hydroxymethylcytosine (5hmC). DNA hypomethylation continues upon liver metastasis. Here we examine 5hmC changes upon progression to liver metastasis.

**Results:** 5hmC is increased in metastatic liver tissue relative to the primary colon tumour and expression of TET2 and TET3 is negatively correlated with risk for metastasis in patients with CRC. Genes associated with increased 5-hydroxymethylcytosine show KEGG enrichment for adherens junctions, cytoskeleton and cell migration around a core cadherin (CDH2) network. Overall, the 5-hydroxymethylcyosine profile in the liver metastasis is similar to normal colon appearing to recover at many loci where it was originally present in normal colon and then spreading to adjacent sites. The underlying sequences at the recover and spread regions are enriched for SALL4, ZNF770, ZNF121 and PAX5 transcription factor binding sites. Finally, we show in a zebrafish migration assay using SW480 CRISPR-engineered TET knockout and rescue cells that reduced TET expression leads to a reduced migration frequency.

**Conclusion:** Together these results suggest a biphasic trajectory for 5-hydroxymethyation dynamics that has bearing on potential therapeutic interventions aimed at manipulating 5-hydroxymethylcytosine levels.

## Background

Colorectal cancer (CRC) is the fourth most common malignancy worldwide[1]. Most deaths in CRC, are because of metastatic disease that occurs in 20% of CRC patients, the liver being the most common site for metastasis (70% of all cases), [2]. DNA methylation changes have emerged as a key driver of metastasis and may explain the organ-specific tropism exhibited by many cancers [3–5]. In CRC, changes in DNA methylation occur alongside genetic alterations during tumorigenesis. These include genome-wide hypomethylation and hypermethylation at specific gene promoters (reviewed [6]). Patient stratification on the basis of epigenetic signatures such as CpG island methylator phenotype (CIMP))[7] [8], and other methylation biomarkers (*VIM*, *SFRP2* and *SEPT9* [9]) are becoming promising therapeutic avenues for targeting CRC [10].

Methylated DNA can be oxidised to form 5-hydroxymethylcytosine (5hmC), that can be further oxidised to 5-formylcytosine (5fC) and 5-carboxylcytosine (5caC) through the action of the Ten-Eleven-Translocation (TET) dioxygenases known as TET1, TET2 and TET3 [11, 12]. These oxidised modifications of 5-methylcytosine are intermediates of active cytosine demethylation and targets for base excision repair. However,5hmC and 5fC have been established as stable epigenetic marks that can be maintained during cell division [13, 14]. Genome-wide sequencing of 5hmC in various mammalian tissues support its role as an active epigenetic marker at enhancers, gene-bodies, and promoters [15, 16]. Using deep neural network models incorporating gene expression and chromatin conformation as well as 5hmC profiles, predictive models of gene expression and putative regulatory regions has recently been developed based on 5hmC signals [17].

Reduced levels of 5hmC have been observed in many cancers in humans [18–21] and mouse cancer models [22, 23]. During normal development, 5hmC and the TETs have a role in cancer-relevant processes including differentiation and stem cell regulation [24, 25]. In a series of CRC patients, we observed that 5hmC levels are globally depleted in adenomas and adenocarcinomas, compared to normal colon, despite the presence of steady state TET mRNA transcripts at all cancer stages [21]. In normal colon, differentiated colonocytes had high levels of 5hmC, whereas stem cells in the crypts had low levels of 5hmC, similar to what we see in the colon tumours [21, 22]. hMeDIP sequencing of normal colon tissue showed that promoters enriched for 5hmC were less likely to become hypermethylated in cancer, despite the loss of 5hmC at these loci in colon tumours [21, 26, 27].

Here we report that the metastatic tumours have higher levels of 5hmC than the primary CRC tumours and that the genome-wide 5hmC profiles are similar to normal colon, suggesting that 5hmC is recovered at specific sites during metastasis. We note that several sites where 5hmC has been recovered, the adjacent CpGs are also hydroxymethylated. The loci where 5hmC recovered and spread were enriched for zink finger transcription factor binding sites SALL4, PAX5, ZNF770 and ZNF121. Integration of 5hmC data with published RNA-seq data [26] showed an enrichment for genes involved in adherens junction and cell migration. Finally, we established that the TETs are important for tumour cell migration using CRISPR-Cas9 mediated triple TET knockout of SW480 cells in a zebrafish xenograft assay.

## Results

### 5hmC levels are increased in liver metastasis tissues compared to primary colon tumours in colon cancer patients

We previously reported reduced 5hmC levels in adenomas and adenocarcinomas [21]. This same cohort included patients (n=10) that at first diagnosis, prior to chemotherapy had liver metastases (Table1). We examined the global 5hmC levels in the liver metastases for comparison to primary tumours in these patients using mass spectrometry and observed that although the global 5hmC levels in liver metastasis samples were reduced compared to normal colon tissue, they were significantly increased compared to the primary carcinoma tissue (Figure 1A).

**Figure 1:**
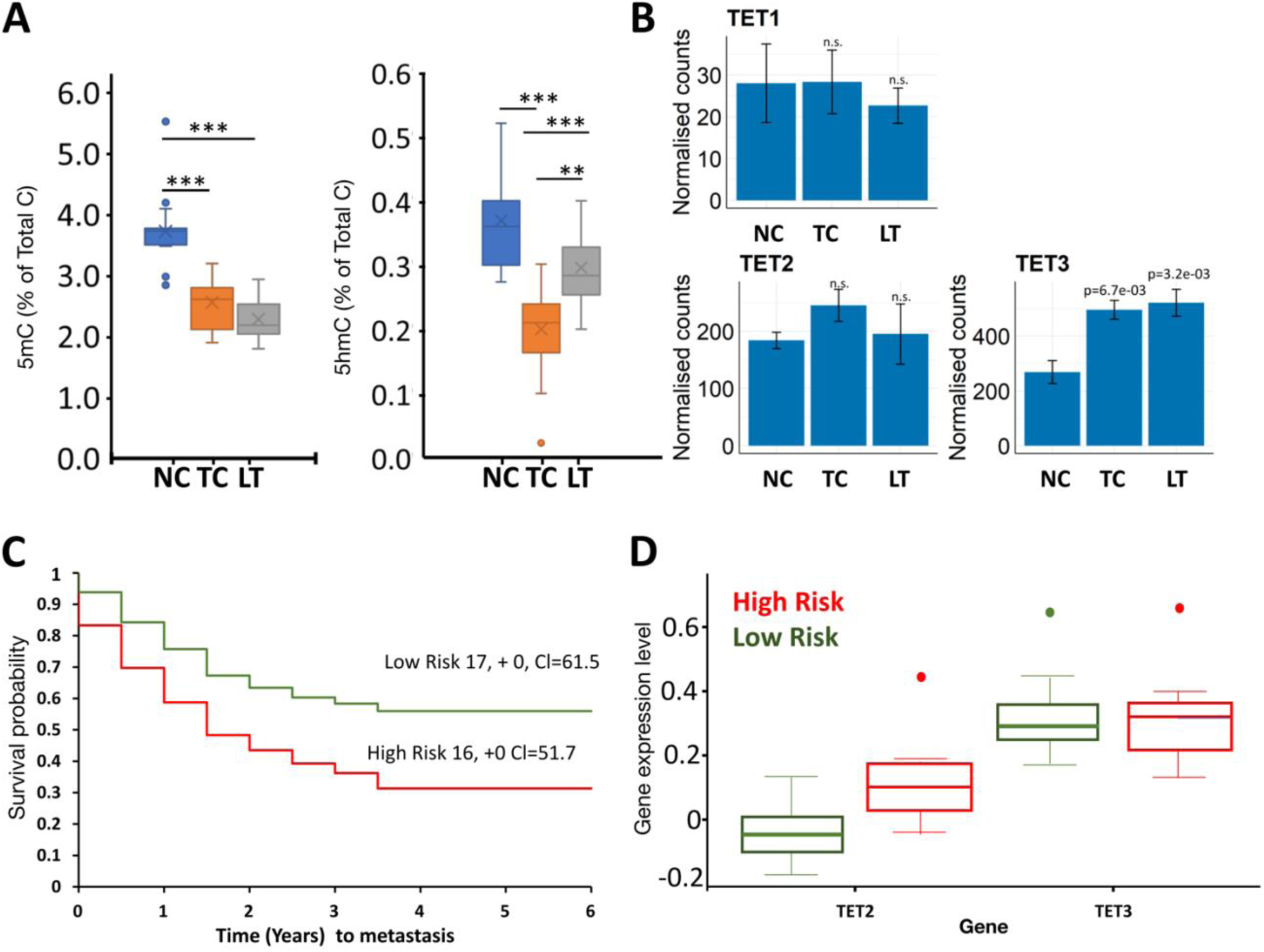
5hmC levels are increased in liver metastasis tissue compared to primary colon tumours In CRC: A) Mass Spectrometer analysis for global 5-methylcytosine (5mC) and 5-hydroxymethylcytosine (5hmC) levels in DNA from 14 normal colon (NC) tissues from CRC patients; 15 tumours in colon (TC) and 10 metastases to liver tumours (LT), showing global demethylation with CRC progression for 5mC. By contrast global 5hmC in metastasis increases compared to the primary cancers. Boxplots depict median and interquartile ranges and standard deviation. *** P< 0.005. B) RNA-seq data from NCBI SRR2089755, showing normalised read counts for TET transcripts in normal colon (NC), tumour colon (TC) and metastasised to liver tumours metastasis (LT). C) Kaplan-Meier plots for time (years) to metastasis generated from the Loboda Yeatman study of Colon cancer (GSE28722), dividing patients into high or low risk for metastasis depending on TET2/TET3 expression. Concordance Index = 57.87, Log−Rank Equal Curves p=0.1093, R^2=0.166/0.994. Risk Groups Hazard Ratio =1.83 (conf. int 0.86-3.86). D) TET2 and TET 3 expression in high and low metastasis risk groups. Patients with high risk of metastasis have higher expression of TET (P=0.000333); no significant difference for TET3expression (P=0.965127).

In our patient cohort the absolute levels of *TET* transcripts were previously measured by standard curve method in adenomas and adenocarcinomas. Despite the reduction in 5hmC at these stages, we found all three *TET*s to have the same mean expression levels as the matched normal tissue, with *TET2* and *TET3* being the most abundant, albeit that there was considerable variation around the mean [21]. No mutations in *IDH1/2* or other genes encoding enzymes known to generate metabolites that inhibit TET catalytic activity were found [21]. We do not have RNA from our metastasis samples, but a publicly available RNAseq dataset for colon cancer with matched liver metastasis [26], confirmed *TET2* and *TET3* to be the more abundant *TET* transcripts in colon and liver metastasis. Unlike our samples, in this dataset, *TET3* expression was significantly increased in colon and liver tumours compared to the normal colon tissue (Figure 1B). Kaplan Meier plots generated using SurvExpress tool8 and colon cancer expression databases (GSE28722) indicated that ‘low-TET2/3’ patients have a better metastasis-free long-term survival (>5years) than ‘high-*TET2/3* (Figure 1C, Concordance Index = 57.87, Log−Rank Equal Curves p=0.1093, R^2^=0.166/0.994). High risk of metastasis was associated with significantly higher *TET2* expression (P=0.0003), while the difference for *TET3* expression between high and low risk patients was not statistically significant (Figure 1D). The overall poor prognosis associated with higher TET expression and the increase in overall 5hmC levels seen in the metastatic samples are counterintuitive to the perception that reduced 5hmC is a hallmark of cancer. We therefore set out to identify the genes that contain 5hmC in the metastatic tumours. We had metastasis samples and matched non-tumour liver tissue for 5 patients that we previously profiled 5hmC in normal and primary carcinoma tissues, using hMeDIP-seq. The same hMeDIP-seq protocol was used to examine 5hmC in the liver metastasis for direct comparison to the primary tumours in these patients. Important questions that we aimed to explore was whether the 5hmC profile in liver metastasis reflects the colon tissue origin of the primary tumour or the metastatic niche, and whether the gain of 5hmC marked metastatic progression.

### 5hmC profiles in liver metastasis match colon profiles

Total counts of hMeDIPseq peaks (enriched over input) for normal colon (NC), primary tumour in the colon (TC), and metastatic tumours to the liver (LT) showed more 5hmC enrichment in liver metastasis compared to primary tumours (Additional File 1-Fig.S1). In normal colon, over 30,000 5hmC peak counts overlapped with about 1% of all cytosines, and 2% of all CpG sites. These were substantially decreased in primary colon tumours (< 800 peak counts, overlapping 0.01% cytosines, and 0.03% CpG sites), and comparatively increased in liver metastasis (1794 peak counts, 0.05% cytosines, 0.1% CpG sites (Additional File1-Fig.S1A). Normal liver (NL) had a higher amount of 5hmC compared to the tumour samples (>20,000 peak counts, 0.5% cytosines, and 1% of CpG sites, Additional File1-Fig.S1A). The distribution of the 5hmC peaks across genomic features showed most peaks to be present at promoters, and the 5’UTR for normal colon and the liver tumours. The colon and liver metastasis tumours followed the same trend with lower overall peak counts (Additional File1-FigS1B). Within genomic regions, the 5hmC peaks followed a profile of highest concentration around the transcription start site (TSS), but an absence at the TSS itself (Additional File1-Fig.S1C). This trend was the same for all tissues, but with more noise in the tumours compared to the normal tissue, due to lower peak numbers.

Given the low amount of 5hmC in tumours, even a small amount of normal liver cells within the metastatic tumours, could increase the overall levels of 5hmC. We analysed our liver metastasis tumours for tumour purity (i.e. fraction of tumour cells), using the AITAC (Accurate Inference of Tumour purity and Absolute Copy number) approach [28], using the input sequencing samples of our normal liver and metastases and found the AITAC tumour purity values to be between 0.84 and 0.86 for the liver metastasis samples (Table 1), which is considered “high purity”.

**Table 1:** Patient samples used for this study.

| CRC Case | Dukes Stage | Available samples |  |  |  | AITAC | sex | Age | Mass Spec | hMeDIP Seq |
| --- | --- | --- | --- | --- | --- | --- | --- | --- | --- | --- |
|  |  | NC | TC | LT | NL | NL |  |  |  |  |
| 37 | C1 | y | y | n | n |  | male |  | y | n |
| 44 |  | y | y | n | n |  | male |  | y | n |
| 51 | D | y | y | y | y | 0.86 |  |  | y | y |
| 52 | D | y | y | y | y | 0.84 |  |  | y | y |
| 53 | D | y | y | y | y | 0.83 |  |  | y | y |
| 55 | D | n | n | y | y | 0.84 |  |  | y | y |
| 56 | C2 | y | y | y | n |  | female | Median Age 73 (range 57 – 81) | y | n |
| 57 | D | y | y | y | n |  |  |  | y | n |
| 58 | C2 | y | y | y | n |  |  |  | y | n |
| 59 | C1 | y | y | y | n |  | female |  | y | n |
| 60 |  | y | y | y | n |  |  |  | y | n |
| 49 | D | n | y | y | y | 0.84 | male |  | y | y |
| 45 |  | y | y | n | n |  | female |  | y | n |
| 48 |  | y | y | n | n |  | female |  | y | n |
| 24 |  | y | y | n | n |  | male |  | y | n |
| 35 | C1 | y | y | n | n |  | male |  | y | n |
| N=16 |  | N=14 | N=15 | N=10 | N=5 | All cases MSI negative, |  |  |  |  |

y=yes, n=no. blank cells information not available, AITAC is a measure of tumour purity.

Principal component analysis (PCA) for 5hmC peaks in metastasis, primary tumour, and normal tissues (Figure 2A), indicated distinct clusters of 5hmC profiles for each phenotype. Normal colon and colon tumours clustered in one dimension, while the normal liver samples mostly clustered away from these, except for one outlier. The liver metastasis samples did not form a tight cluster, with three out of the five samples tending towards the colon clusters and the other two samples tending towards the normal liver cluster (Figure 2A). The intersection of peaks is shown in an upset plot in figure 2B. As expected, normal liver and normal colon have the most 5hmC loci (57.9% of these are unique to liver, and 28.2% unique to colon), and the largest intersection is the 7.1% shared peaks between normal liver and colon (Figure 2B). Considering that the set size for normal liver is 1.7 times higher than for normal colon, we’d expect random overlap between normal liver and liver metastasis to be higher than an overlap with normal colon. However, even though liver metastasis tumours shared 5hmC peaks with both normal colon and with normal liver (1.7%), the intersection with normal liver only was 1.3%, compared to normal colon which was 1.1%.

**Figure 2:**
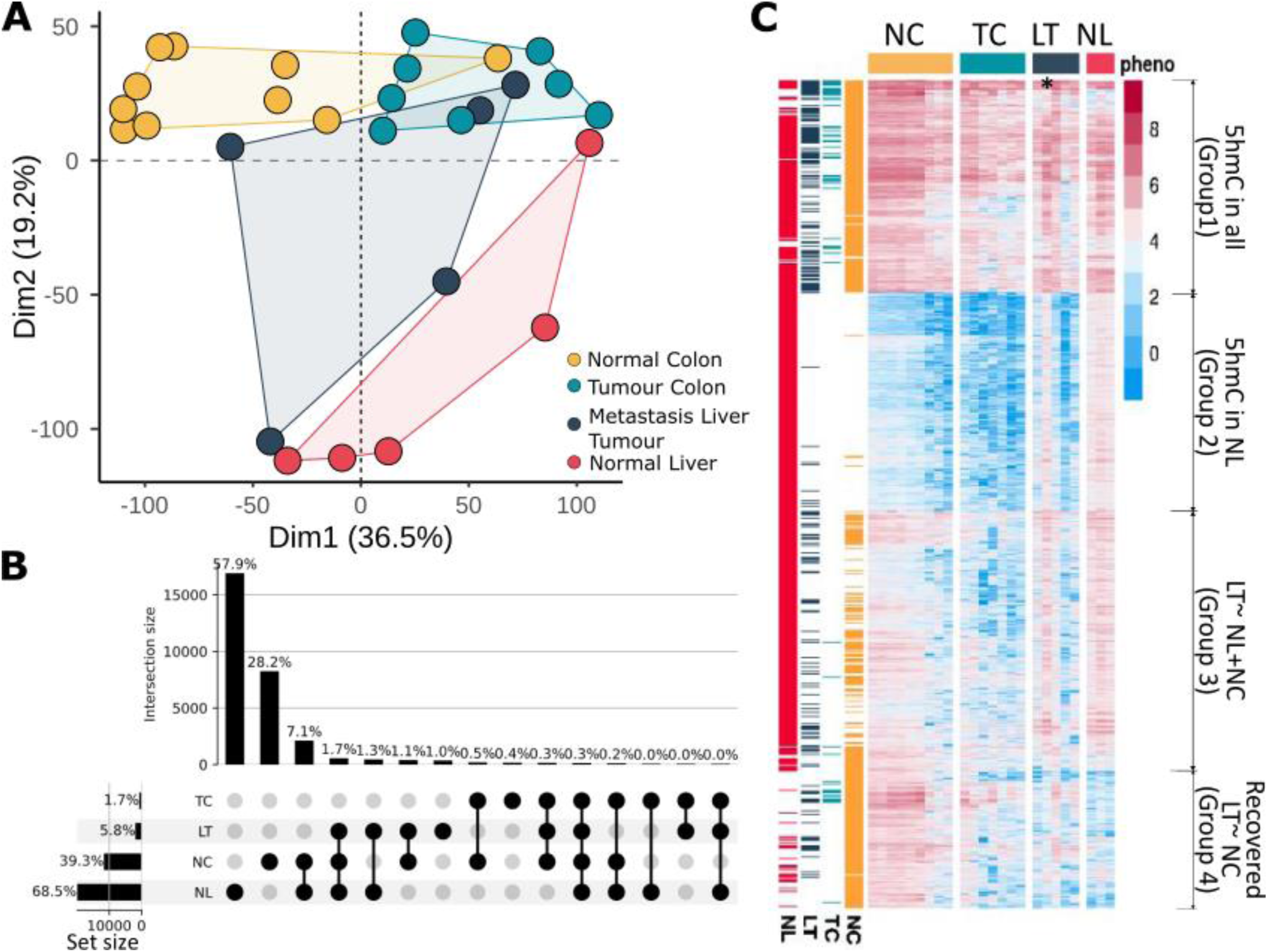
5hmC profiles in liver metastasis and colon are similar. Colour scheme: yellow normal colon tissue from CRC patients (NC); light blue primary tumour in colon (TC); dark blue metastasized to liver tumours (LT); red normal liver (NL). A) Principal Component Analysis plot showing distinct clustering between NC (n=9), TC (n=8), NL (n=5), and LT (n=5). B) Upset plot showing the overlap of 5hmC peaks in normal and tumour tissues. C) Heatmap of 5hmC profiles during CRC tumour progression; first four columns depict average 5hmC levels in each of NL, LM, CT, NC. Subsequent columns depict individual patient samples grouped by tissue type. Four distinct clusters are present: Group 1) 5hmC presence in all samples; Group 2) 5hmC present in NL only; Group 3) liver metastasis with similarities to both NC and NL; and Group 4) recovery of 5hmC in LT matching NC. Star * highlights patient sample LT51, which looks very similar to normal liver.

A heatmap analysis to examine differential 5hmC levels between samples (Figure 2C), highlighted distinct 5hmC profiles in normal liver and colon. Colon tumours had similar clustering to normal colon, albeit fewer loci with 5hmC. The liver metastasis samples had loci that fell within 4 hierarchical groups, “group 1” having high 5hmC common to normal colon and liver; “group 2” low 5hmC similar to normal colon and colon tumours; “group 3” reduced levels of 5hmC compared to normal tissue, but more 5hmC than colon tumours; and “group 4” showing a cluster of genes that appear to have gained 5hmC in the liver metastasis samples at the same loci as in the normal colon (Figure 2C). One exception was a patient in which the liver metastasis had a similar 5hmC profile to normal liver. This metastasis sample is the one that clustered with the normal liver in the PCA, possibly suggesting contamination from adjacent normal liver at the time of the biopsy. We repeated the PCA and heatmap analyses without this potentially contaminated sample. This resulted in a tighter cluster of liver metastasis in the PCA (Additional File1-Fig. S1D). With or without the inclusion of this sample (Additional File1-Fig. S1E,F), the heatmaps still showed a clear cluster of loci that have 5hmC in the liver metastasis, shared with the normal colon and to a lesser extent the colon tumours, and mostly absent in the normal liver. Interestingly, this potentially contaminated sample had a high purity AITAC score, suggesting that in addition to CNV, other parameters (tumour aneuploidy, SNPs and heterogeneity) and possibly even 5hmC profiles should be considered in these pipelines to provide a more accurate estimation of tumour purity.

### A 5hmC signature in liver metastasis is associated with the activation of genes associated with adherens junctions, regulation of cytoskeleton, calcium dependent and independent cell matrix adhesion and focal adhesion

Additional File 2-Table S1 lists the genomic coordinates for consensus peaks for the different CRC cancer stages. We examined the associated genes that had 5hmC peaks within the gene body and 5kb upstream of the TSS. KEGG pathway analysis for 1548 genes associated with consensus 5hmC peaks uniquely present in the metastatic samples showed significant enrichment for genes involved in viral and bacterial infection indicative of characteristic CRC microbial dysbiosis, as well as metastasis related functions such as adherens junction and focal adhesion (Figure 3A).

**Figure 3:**
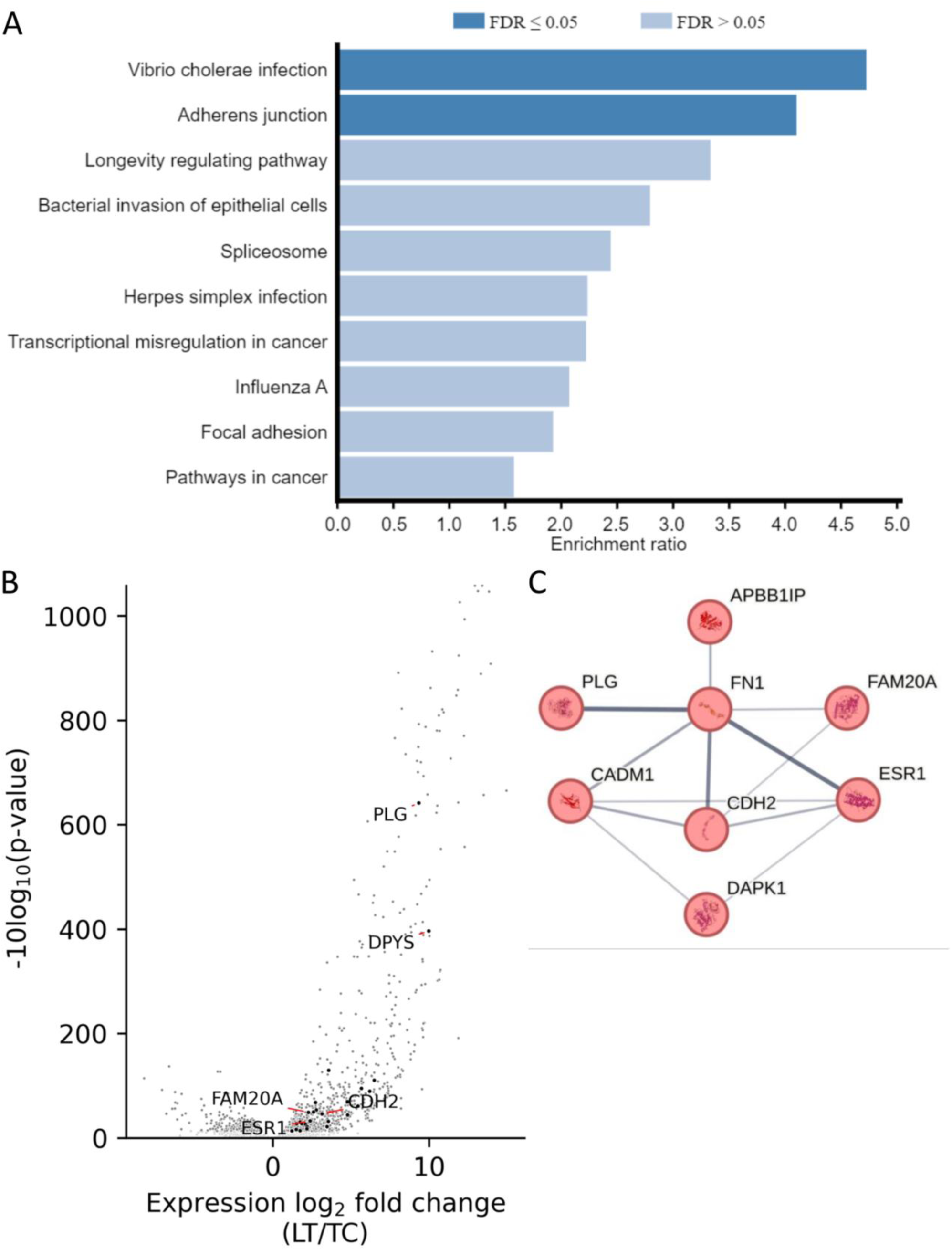
5hmC peaks associated with differential gene expression. A) KEGG analysis of 5hmC marked genes that gain 5hmC in liver metastasis B) Volcano plot for gene expression changes after metastatic transition (data from NCBI SRR2089755). Darker dots identify genes with at least one overlapping and significantly enriched 5hmC peak in hMeDIP-seq. C) A core Cadherin 2 (*CDH2*), and Fibrinogen (*FN1*) network of associations for genes that have increased 5hmC and upregulated expression during metastasis.

Repeating the KEGG analysis with the genes associated with 5hmC changes between primary and metastatic tumour returned enrichment for adherens junction, regulation of cytoskeleton, and focal adhesion. To determine whether these 5hmC changes constitute a metastasis signature linked with gene expression, we used RNAseq data [26] and integrated 5hmC changes with differential gene expression (Additional File 3 -Tables S2). Figure 3B illustrates differential gene expression between the liver metastasis and the primary colon tumours, with 5hmC marked genes superimposed. In liver metastasis, 2% (365) of the 20470 transcripts identified (the whole transcriptome), were associated with genes that have 5hmC peaks overlapping the transcription unit or promoter. Most of tshese (324 5hmC marked loci) were not associated with the 2891differentially expressed genes (−0.95 > *log*_2_*FC* or *log*_2_*FC* > 0.95). Out of the 934 transcripts that were down regulated, 5(1%) were marked by 5hmC, compared to 37(2%) of the 1957 transcripts that were upregulated. Thus out of the 5hmC marked genes, 42 (12%) were associated with differential expression, of which 37 (88%) were upregulated and 5 (12%) were downregulated in metastasis (Additional File 4 – Table S3). The most highly expressed gene on this list was *PLG* (Plasmin heavy chain A), which had a 28-fold increase in expression in liver metastasis compared to the primary tumour. We submitted the 37 upregulated with 5hmC to STRING analysis with k-means clustering (Additional File1-FigS2), which returned a higher-than-expected number of interactions for genes involved in metastasis pathways including calcium dependent and independent cell matrix adhesion, wound healing, and migration. A cluster with the most high-confidence edges included cadherin 2 (*CDH2*), and fibronectin (*FN1*) (PPI enrichment P-value 9.31e-10) (Figure 3C). These two proteins are involved in cell adhesion and migration, hence their connection in the STRING. While they operate through different mechanisms, and serve distinct functions, it is likely that they indirectly influence each other during metastatic processes and that the genes in this cluster (Table 2) which also include *PLG* and *ESR1* constitute a 5hmC metastatic signature for colon cancer.

**Table 2:**
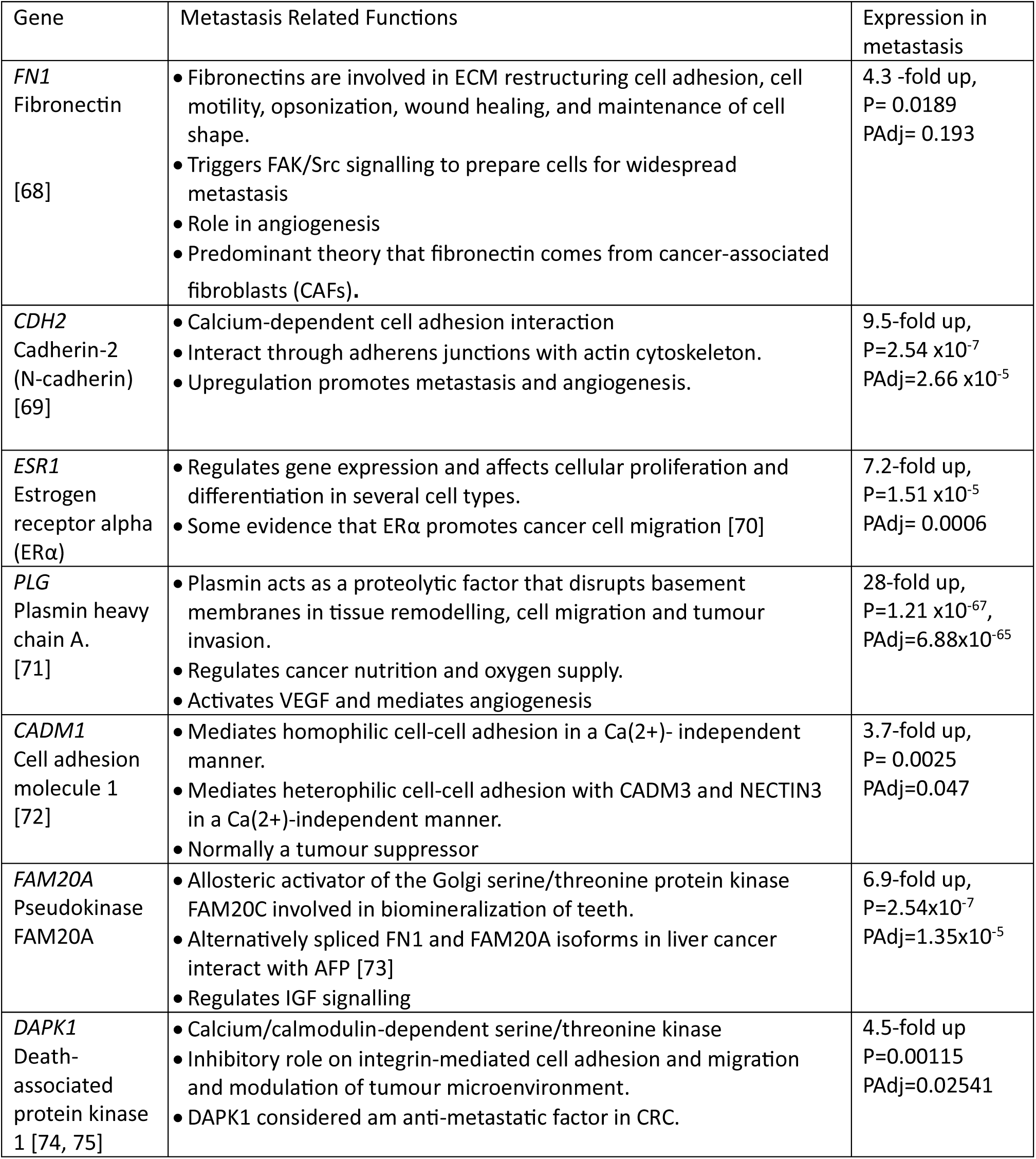
Network of genes with 5hmC and upregulated transcription in metastatic tissue.

| Gene | Metastasis Related Functions | Expression in metastasis |
| --- | --- | --- |
| *FN1*<br>Fibronectin<br>[68] | • Fibronectins are involved in ECM restructuring cell adhesion, cell motility, opsonization, wound healing, and maintenance of cell shape.<br>• Triggers FAK/Src signalling to prepare cells for widespread metastasis<br>• Role in angiogenesis<br>• Predominant theory that fibronectin comes from cancer-associated fibroblasts (CAFs). | 4.3 -fold up,<br>P= 0.0189<br>PAdj= 0.193 |
| *CDH2*<br>Cadherin-2<br>(N-cadherin)<br>[69] | • Calcium-dependent cell adhesion interaction<br>• Interact through adherens junctions with actin cytoskeleton.<br>• Upregulation promotes metastasis and angiogenesis. | 9.5-fold up,<br>P=2.54 x10<sup>-7</sup><br>PAdj=2.66 x10<sup>-5</sup> |
| *ESR1*<br>Estrogen<br>receptor alpha<br>(ERα) | • Regulates gene expression and affects cellular proliferation and differentiation in several cell types.<br>• Some evidence that ERα promotes cancer cell migration [70] | 7.2-fold up,<br>P=1.51 x10<sup>-5</sup><br>PAdj= 0.0006 |
| *PLG*<br>Plasmin heavy<br>chain A.<br>[71] | • Plasmin acts as a proteolytic factor that disrupts basement membranes in tissue remodelling, cell migration and tumour invasion.<br>• Regulates cancer nutrition and oxygen supply.<br>• Activates VEGF and mediates angiogenesis | 28-fold up,<br>P=1.21 x10<sup>-67</sup>,<br>PAdj=6.88x10<sup>-65</sup> |
| *CADM1*<br>Cell adhesion<br>molecule 1<br>[72] | • Mediates homophilic cell-cell adhesion in a Ca(2+)- independent manner.<br>• Mediates heterophilic cell-cell adhesion with CADM3 and NECTIN3 in a Ca(2+)-independent manner.<br>• Normally a tumour suppressor | 3.7-fold up,<br>P= 0.0025<br>PAdj=0.047 |
| *FAM20A*<br>Pseudokinase<br>FAM20A | • Allosteric activator of the Golgi serine/threonine protein kinase FAM20C involved in biomineralization of teeth.<br>• Alternatively spliced FN1 and FAM20A isoforms in liver cancer interact with AFP [73]<br>• Regulates IGF signalling | 6.9-fold up,<br>P=2.54x10<sup>-7</sup><br>PAdj=1.35x10<sup>-5</sup> |
| *DAPK1*<br>Death-<br>associated<br>protein kinase<br>1 [74, 75] | • Calcium/calmodulin-dependent serine/threonine kinase<br>• Inhibitory role on integrin-mediated cell adhesion and migration and modulation of tumour microenvironment.<br>• DAPK1 considered an anti-metastatic factor in CRC. | 4.5-fold up<br>P=0.00115<br>PAdj=0.02541 |

It has been demonstrated that the DNA hypermethylation profile in primary colon is maintained when cells migrate to the liver [27]. We previously reported that 5hmC-marked promoters from normal colon are protected from hypermethylation in primary tumours [21]. In contrast, gene promoters that had gained 5hmC in colon tumours had loss of DNA methylation [21]. Since methylated DNA is a substrate for TETs, we re-examined the colon tumour methylation data from our previous study to see whether the above panel of 5hmC metastasis signature genes, were methylated. *CDH2* and *ESR1,* two of the key genes in the 5hmC metastasis signature were significantly hypermethylated at their promoters (P<0.0001) in colon cancer compared to normal colon. In a published MBD-seq dataset [27] for colon tumour and liver metastasis tissue, we confirmed that *CDH2* and *ESR1* were hypermethylated in primary colon cancer consistent with our findings. In this dataset 88% of genes that gained 5hmC in the liver metastasis were methylated in the primary tumours, which is 10% more than the overall percentage of genes that had methylation in the primary tumours. However, the gain of 5hmC was not accompanied by a reduction in DNA methylation in the metastasis tumours (Additional File 3-table S2). The overlap of 5mC and 5hmC in metastasis, could suggest a dynamic turn-over between methylation and demethylation but more likely reflects the heterogeneity of tumours with different subsets of cells having either methylation or hydroxymethylation. A further consideration is that the 5hmeDIP-seq technique enriches for 5hmC using single stranded DNA as input, whereas MBD-seq uses double stranded DNA [29]. Although MBD is highly specific for 5mC and is inhibited by 5hmC [30], it is feasible that some loci could be asymmetrically modified with 5mC and 5hmC on different strands of the same DNA molecule.

### 5hmC “recovery” spreads to adjacent CpGs at a subset of loci

A visual examination of the 5hmC data on an integrative genomics viewer (IGV), showed the expected pattern of 5hmC “recovering” at sites where 5hmC was present in normal colon tissue, absent in the colon tumour, and recovered in the metastatic tumours (Additional File1-FigS3). Interestingly, we observed several regions, where the 5hmC had not only recovered, but spread to form adjacent peaks that were not present in the normal tissue. Figure 4A is an example of the *LIMS2* locus showing both recovered and spreading of 5hmC. To determine how frequently the “recovery and spreading” occurred, we relaxed the stringency for consensus peaks to 50% and merged the 5hmC peaks that were close to each other (within 500bp) in the liver metastasis data to obtain a list of merged peaks that had spread (n=15939, a minimum size of 1000bp) and refined the list for overlap between normal colon (n=3149, Figure 4b, see methods for detail). Since 5hmC is high in normal liver, we ran a similar merging of peaks in normal liver data and obtained 2328 merged and spread peaks, that we compared to the “merged in metastasis” peaks. This left 2405 unique “RS-peaks” in the liver metastasis where 5hmC had Recovered after an absence in the primary tumour and then Spread to adjacent sites. We then randomly shuffled the NC, TC and LT merged peaks 1000 times and found that on average 522 recover and spread peaks were observed. A permutation test revealed that the 3149 recover and spread peaks we observed occur significantly more than expected than by chance (p < 0.001) and shows that the RS-peaks that we found are not occurring randomly.

**Figure 4:**
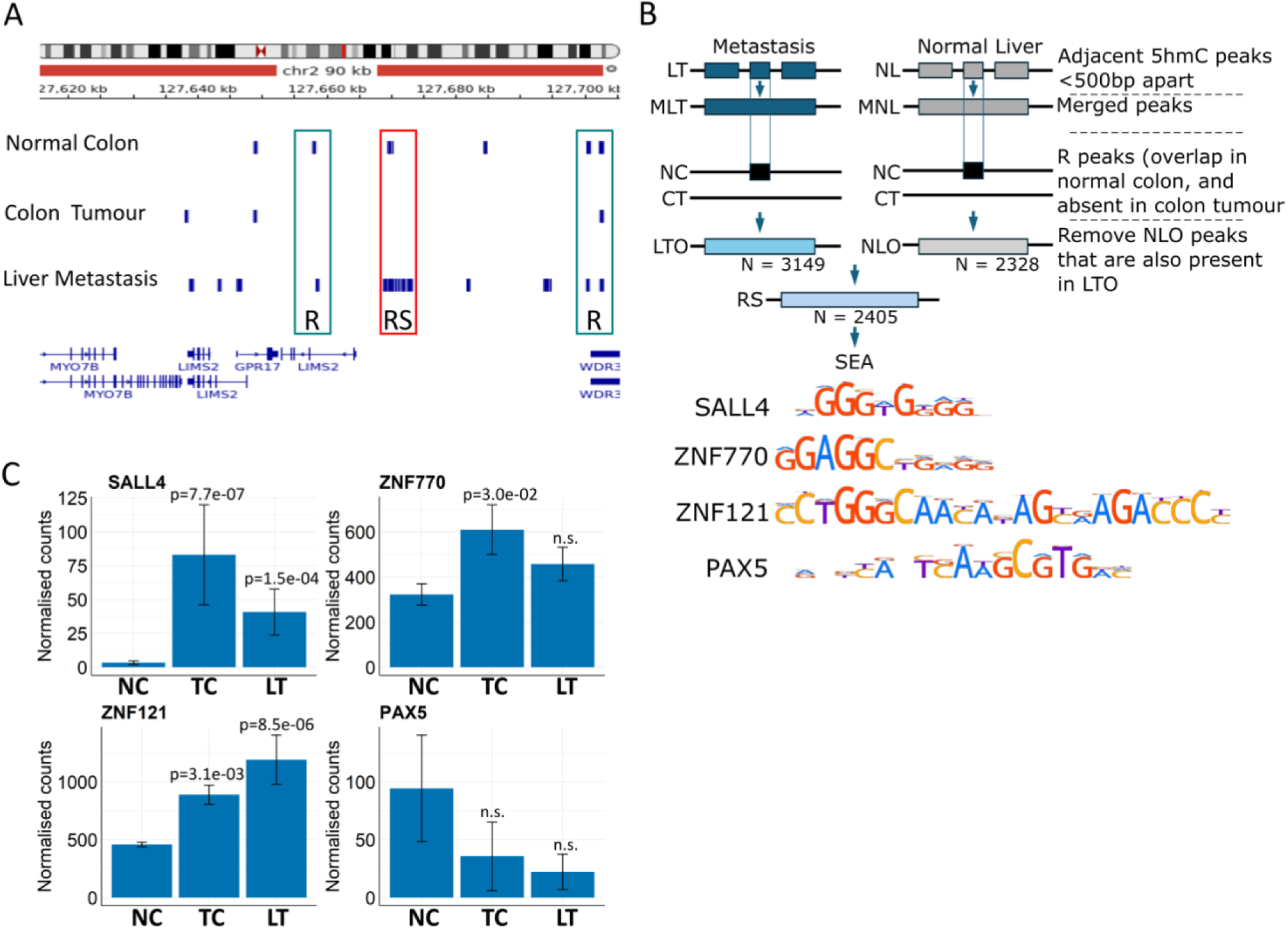
In liver metastasis 5hmC peaks that were initially present in normal colon are recovered. A) IGV screenshot with uploaded f 5hmC consensus peaks for normal colon, colon tumour and liver metastasis. Boxes above R-highlight a 5hmC peak that has “recovered” since it is present in normal colon, absent in colon tumours and reappears in liver metastasis. Boxes above RS-highlight a 5hmC peak that has “recovered and spread”. Thus, the peak is present in normal colon, lost in tumour colons and reappears in metastasis, but also spreads to adjacent sites. B) The strategy for identifying RS peaks and the consensus binding sites for SALL4, ZNF770, ZNF121 and PAX5 identified after SEA. Abbreviations: MLT and MNL are **M**erged liver tumour (LT) and normal liver (NL) peaks respectively. LTO and NLO are LT or NL peaks that **O**verlap normal colon. C) Normalised RNAseq counts (NCBI SRR2089755) for *SALL4*, *ZNF770*, *ZNF121*, and *PAX5* in normal colon, colon tumours and liver metastasis (NC, CT and LM). Wald test, Benjamini-Hochberg adjusted. Error bars represent ± SEM.

The underlying sequences were then examined for enrichment of specific transcription factor binding sites. The most frequently enriched transcription factor binding sites included SALL4 (n= 6879, 90.4%), PAX5 (n=5345, 70.2%), ZNF770 (n= 4363, 57.3%) and ZNF121 (n= 3843, 50.4%) (Figure 4B). For these the total number of binding sites were: SALL4: 11825 (1.71 per sequence); ZNF770: 8414 (1.92 per sequence); PAX5: 7665 (1.43 per sequence) and ZNF121 = 3851 (1.00 per sequence). The coordinates for these “recovered and spread” 5hmC peaks are listed in Additional File 5-Table S4.

*PAX5* transcripts are present at very low levels in normal tissue and are further reduced in tumours, whereas *SALL4, ZNF770* and *ZNF121* transcripts are upregulated in primary and metastatic tumours compared to normal colon (Figure 4C). Liver metastasis tends to have lower expression of *SALL4* and *ZNF770* compared to the primary tumours (Figure 4C). Thus, *SALL4* and *ZNF770* transcription was inversely correlated with 5hmC levels during progression from normal to primary colon cancer to liver metastasis.

### TET expression in a colon cancer cell line SW480 increases the incidence of cell migration in Zebrafish xenograft assays

The 5hmC signature included a core of genes associated with cell migration (*CDH2, FN, ESR1*), suggesting that the TETs have a role in metastatic progression. We predicted that targeting 5hmC via the TETs in a colon cancer cell line would inhibit cell migration in a xenograft assay. Embryo-larva-zebrafish are increasingly being shown to be an excellent model for human cell transplantation and cell migration [31–33] and SW480 cells have been shown to successfully implant and migrate in zebrafish larvae [34, 35]. We generated a CRISP-Cas9 SW480 TET1-3 triple knockout cell line (SW480TKO, Additional File 1-Fig4A-E). The SW480TKO cell line had significantly depleted TET mRNA and protein levels (Additional File1-Fig.S4B and C). A TET activity assay confirmed enzymatic activity in the wild type cells and showed substantial reduction in the SW480TKO cells (Additional File1-Fig S4D), consistent with reduced 5hmC levels measured by immunoblotting (Additional File1-Fig.S4E). The low levels of persistent 5hmC in SW580TKO suggest a “knockdown” of TET proteins rather than a complete knock out. The reduction of TETs and 5hmC had no effect on the global DNA methylation levels (Additional File1-FigS4E). No significant differences in proliferation- or colony forming rate was observed between the SW480 wildtype and TKO cells (results not shown).

SW480 wildtype and TKO cells were injected into the perivitelline space of zebrafish larvae. Those that developed a tumour mass in the yolk sac, with or without migration to distant sites, were scored after 48 hours. We limited migration scoring to the tail since this has less auto-fluorescence than sites such as the eye or brain. Examples of tumour mass in the yolk sac and distant foci are shown in Figure 5A. For the SW480 wildtype cells (n=60), a primary tumour mass in the yolk sac was observed in 37 larvae, of which 15 (40%) had evidence of migration with an average number of 14.75 distant foci per animal (Table 3). A total of 160 larvae were injected with the SW480 TKO cells (two clones with 92 and 68 respectively), and primary tumour mass was observed in 137 larvae. Out of these, only 15% of larvae had tumour cells that had migrated away from the primary tumour with an average number of 14 foci per animal (p<0.05 Table 3). Thus, significantly fewer larvae that received the SW480-TKO cells had evidence of cells migrating from the yolk sac to the tail compared to embryos that received the unmodified SW480 cells (Figure 5B). Aside from the significant difference in migration rate between the wild type and TKO cells in the xenograft assay, no differences were seen in the size of tumours, or the number of metastatic foci per fish. To determine whether we could rescue the TKO migration effect, we transiently transfected a mouse *Tet2* construct in the SW480TKO cells before injecting these into zebrafish embryos. The ectopic expression of *Tet2* resulted in increased global 5hmC levels compared to the SW480TKO cells (Figure 5C and D) and showed an increase in migration in the zebrafish assay, with 34% of the larvae having distant foci compared to 18% in untreated SW480TKO cells (Figure 5B). These results support a role for the TET proteins in mesenchymal cell migration.

**Figure 5:**
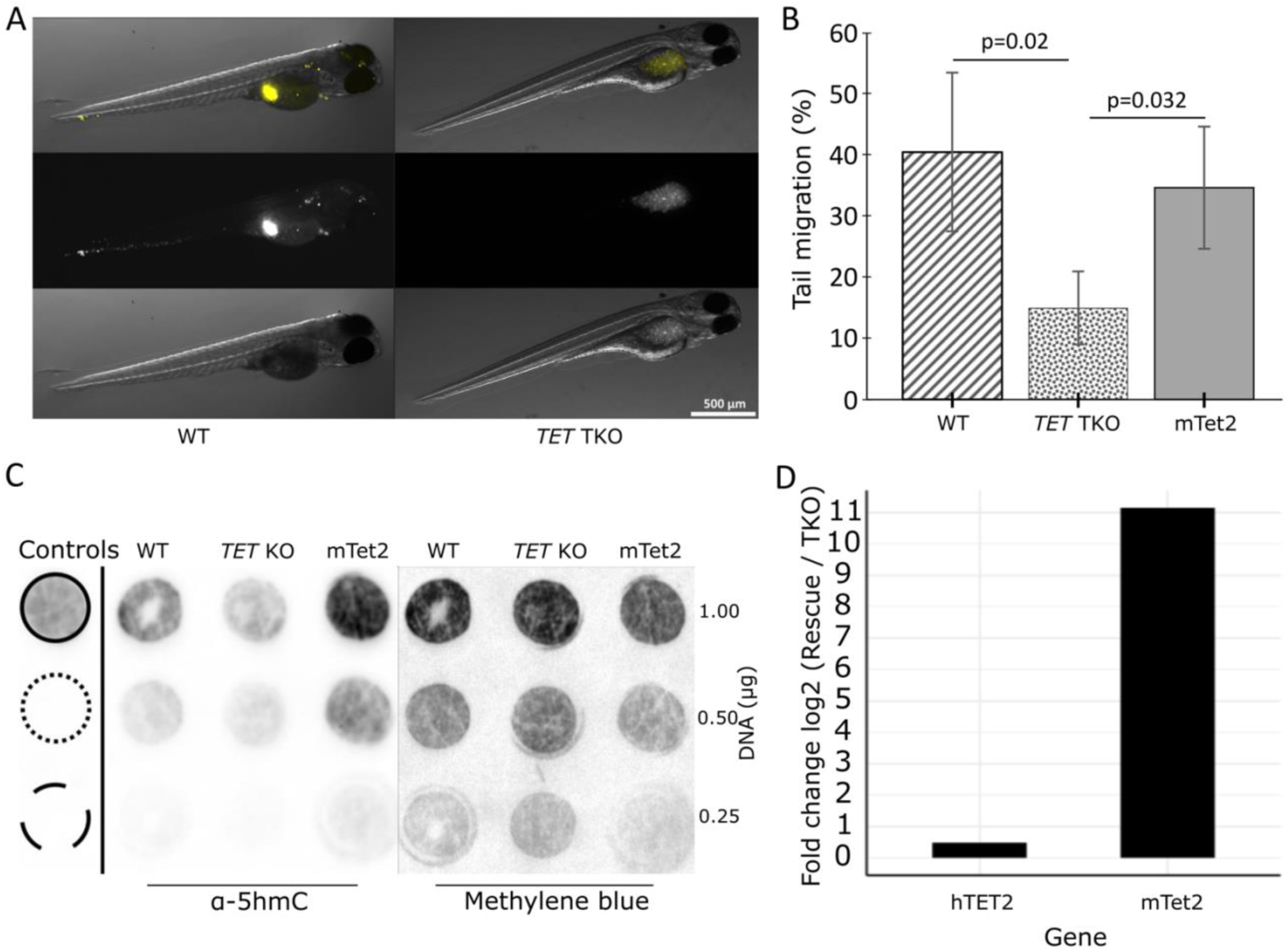
Reducing TET expression and 5hmC levels inhibits migration in a zebrafish xenograft assay. A). Bright field and fluorescence examples of two larvae with accumulation of DIL stained SW480 cells within the perivitelline space and the yolk sac. The panels on the left show cells that have migrated towards the head and tail, the panels on the right show cells remaining at site of injection and no migration. B). Incidence of cell migration (number embryos with foci in tail/all embryos with cells in yolk sac), after injecting SW480 cells (WT), CRISPR *TET1-3* (TKO) and transiently expressed mouse *Tet2* (TKO + m*Tet2*), also see Table 3 for statistics. C) A representative immunoblot for 5hmC levels in WT, TKO and *mTet2* rescue cells, confirming that 5hmC is reduced in TKO and rescued with transient m*Tet2* expression. Antibody controls include spots for nucleotide standards: 5hmC top, 5mC middle and unmodified C bottom. D) Representative RT-qPCR assay confirming ectopic expression of mouse (m*TeT2)* in the TKO cells after transient transfection. As a control we used a PCR assay for human (hTET2) which measures residual endogenous *TET2* expression.

**Table 3:** Zebrafish larvae Migration assay.

|  | Embryos injected | Embryos with Tumour at injection site (%) | Embryos with Migration | Migration Incidence (%) | Number of Foci | Average Number of Foci |
| --- | --- | --- | --- | --- | --- | --- |
| TKO vs WT |  |  |  |  |  |  |
| SW480 WT<br>4 injection cycles<br>Average<br>SD | 6 | 6 (100) | 3 | 50.00 | 8;12;7 | 9.00 |
|  | 15 | 5 (33) | 1 | 20.00 | 14 | 14.00 |
|  | 21 | 13(62) | 5 | 38.46 | 36;12;11;5;10 | 14.80 |
|  | 18 | 13 (72) | 6 | 46.15 | 6;5;46;26;17;8;12 | 17.14 |
|  | 15.00 | 9.25 (67) | 3.75 | 38.65 |  | 13.74 |
|  | 6.48 | 4.35 (27.5) | 2.22 | 13.33 |  | 3.43 |
|  | N=60 | N=37 | N=15 |  |  |  |
| SW480 TET TKO 2I7<br>4 cycles<br>Average<br>SD | 32 | 24 (75) | 5 | 20.83 | 5;13;5;9;18 | 16.67 |
|  | 19 | 13 (68) | 0 | 0.00 |  |  |
|  | 20 | 19 (95) | 3 | 15.79 | 5;21;24 | 16.00 |
|  | 21 | 19 (90.5) | 4 | 21.05 | 15;17;18;14 | 16.33 |
|  | 23.00 | 18.75 (82) | 3.00 | 14.42 |  | 16.33 |
|  | 6.06 | 4.50 (13) | 2.16 | 9.92 |  | 0.33 |
|  | N=92 | N=75 | N=12 | P=0.036* |  | P>0.05 |
| SW480 TET TKO 6B12<br>4 cycles<br>Average<br>SD | 13 | 12 (92) | 2 | 16.67 | 14;7 | 10.50 |
|  | 16 | 15 (94) | 1 | 6.67 | 6 | 6.00 |
|  | 14 | 12 (86) | 2 | 16.67 | 6;26 | 16.00 |
|  | 25 | 23 (92) | 2 | 8.70 | 10;6 | 8.00 |
|  | 17.00 | 15.50 (91) | 1.75 | 12.17 |  | 10.13 |
|  | 5.48 | 5.20 (4) | 0.50 | 5.25 |  | 4.33 |
|  | N=68 | N=62 | N=7 | P=0.01* |  | P>0.05 |
| WT vs ALL TKO | P=0.002* |  |  |  |  |  |
| Rescue mTet2 vs TKO |  |  |  |  |  |  |
| SW480 TKO 2I7<br>3 cycles<br>Average<br>SD | 19 | 8 (42) | 2 | 25.00 | 6;5 | 5.5 |
|  | 23 | 23(100) | 3 | 13.04 | 12;63;24 | 33.0 |
|  | 12 | 12(100) | 3 | 25.00 | 8;18;24 | 16.7 |
|  | 18.00 | 14.33 (81) | 2.67 | 21.01 |  | 18.39 |
|  | 5.57 | 7.77 (33.4) | 0.58 | 6.90 |  | 13.83 |
|  | N=54 | N=43 | N=8 |  |  |  |
| SW480 TKO 2I7 + mTet2<br>3 cycles<br>Average<br>SD | 11 | 6 (55) | 3 | 50.00 | 23;31;52 | 37.66 |
|  | 26 | 26 (100) | 7 | 26.92 | 9;97;5;41;12;15;16 | 27.86 |
|  | 18 | 17 (94) | 7 | 41.18 | 17;7;6;89;75;81;24 | 42.14 |
|  | 18.33 | 16.33 (83) | 5.67 | 39.37 |  | 35.89 |
|  | 7.51 | 10.02 (25) | 2.31 | 11.64 |  | 7.31 |
|  | N=55 | N=49 | N=17 | P=0.032** |  | P>0.05 |

\*Two sample equal variance (TKO vs WT). \*\* Paired T-test for rescue (mTet2 vs TKO).

Altogether, our data suggest that 5hmC levels follow a biphasic trajectory during colon cancer progression to metastasis, where many loci that fail to accumulate 5hmC in primary tumours restore 5hmC in metastasis. TET-mediated regulation of key genes with functions associated with cell migration may facilitate the transition from primary tumour to metastasis. A subset of loci that regain 5hmC also accumulate 5hmC at adjacent CPGs potentially mediated by the Krab-Zn finger transcription factors ZNF770 and ZNF121, which themselves are regulated by TETs.

## Discussion

Global 5hmC levels are distinctively cell-and tissue-specific, as a result of tissue-specific differences in mitotic cell cycling, TET expression and cofactors that influence the turn-over and accumulation of 5hmC within a cell (reviewed [36]). It has been suggested that the distribution of 5hmC could be an “imprint” of cell identity in various normal tissues, acquired during adult progenitor cell differentiation [36]. Cancer-associated 5hmC signatures for colon cancer and other cancer types have been studied in cell free DNA [37]. It has further been shown that 5hmC is reduced compared to normal cells in almost all primary tumours and haematological malignancies (reviewed in [38]) suggesting that restoring 5hmC levels through the upregulation of TET activity could be a therapeutic avenue for cancer treatment [39–42]. The fact that several cofactors including Vitamin C have a positive effect on upregulating TET activity and increasing 5hmC levels [43–48], provides further opportunities for holistic adjuvant treatments. Our study showing that 5hmC levels are increased in metastatic tumours of the liver raise a cautionary caveat for therapeutic approaches aimed at increasing TET activity in colon cancer patients. An independent data set (Loboda Yeatman study of Colon cancer (GSE28722) [49]) demonstrated poor metastasis-free survival in patients with high *TET2* expression levels and our zebrafish xenograft assays further support a role for *TET2* in metastatic transformation in colon cancer. In our patient cohort there was considerable variation in the absolute levels of *TET* transcripts in primary colon tumours and the matched normal tissue [21]. More patients would be required to stratify the risk according to high and low TET expression and to correlate this with genetic polymorphisms and splice variants of TET genes. While our earlier studies, found no correlation with 5hmC levels or any mutations in IDH1/2 or other enzymes involved in generating metabolites that could inhibit TET catalytic activity [21], there is an extensive number of miRNAs that may affect TET2 function, including miR-7, miR-125b, miR-29b/c, miR-26, miR-101, miR142, and Let-7 [50], post transcriptional modifications and numerous interacting transcription factors, chromatin modifiers and signalling proteins (reviewed in [51].

A literature search for examples of poor prognosis and TET expression showed the TETs are predominantly tumour suppressors in most cancers and predict a worse outcome with low TET expression levels. However, several examples of oncogenic tumour promoting instances have been reported in breast [52, 53], gastric [54], lung [55], ovarian cancers [56], hepatocarcinoma [57] and glioma [58]. In breast cancer, hypoxia induced upregulation of *TET1/TET3* transcription and increased 5hmC levels was reported to drive metastatic transformation and poor prognosis [53]. TET1 and TET 2 promoters contain contain consensus binding sites for HIF1-alpha and upon induction by hypoxia, the resultant changes in 5hmC was associated with increased TNFalpha expression and activation of the TNF-alpha-p38-MAPK signalling [53]. Another example is in hepatocarcinoma where increased TET2 expression has been associated with poor prognosis [57]. In that study TET2 was shown to mediate the canonical EMT E-cadherin-N-cadherin switch by repressing the E-cadherin promoter via recruitment of HDAC1 leading to the activation of B-catenin [57]. These examples indicate context dependent oncogenic/tumour suppressing activity for TETs and 5hmC levels. A precedent for such opposing tumour promoting/suppressor functions is the well described case of the TGF-beta gene which acts as a tumour suppressor by inhibiting proliferation and inducing apoptosis in various cell types, and yet the overexpression of TGF-beta in tumour cells induces epithelial-mesenchymal transitions and promotes invasiveness and metastasis [59].

Our study illustrates the difficulty in ascribing either 5hmC or the TET proteins as causative factors in colon cancer metastsis. While a global increase in 5hmC in metastatic tumours may reflect increased TET activity in the nucleus, the non-catalytic functions of TET proteins cannot be excluded. Thus the TETs may transcriptionally activate or repress target genes to drive metastasis, through the interaction with histone deacetylase 2 (Hdac 2), O-glcNAC transferase (OGT), Sin 3A complex and hypoxia-inducible factors (reviewed [51]). The genome-wide profile in this study showed that 5hmC in the metastatic tumours were similar to their cells of origin (colon) and were mostly associated with active gene expression. A limitation of 5hMeDIP methodology is that it although it provides good overall genome coverage, it doesn’t provide base-pair resolution of 5hmC. Newer single cell sequencing technologies will resolve the uncertainties of tumour purity and intertumoral heterogeneity and enable the identification of rare cell populations and clonal evolution within primary and metastatic tumours.

KEGG pathway analysis for the genes nearest and overlapping the 5hmC peaks in the liver metastasis highlighted enrichment for adherens junction, cytoskeleton and cell migration pathways, suggesting that 5hmC remodelling contributes to the metastatic mesenchymal transition. Filtering for the genes with the most significant changes in gene expression delivered a core cluster of genes enriched for metastasis pathways including calcium dependent and independent cell matrix adhesion, and fibroblast migration. This cluster included cadherin 2 (*CDH2*), and fibronectin (*FN1*), both of which are involved in cell adhesion and migration through different mechanisms. Future analyses in a larger cohort of primary colon and liver metastasis samples examining the combined effects of these known biomarkers will determine whether 5hmC has a role in mediating any synergistic effects during metastasis.

Colon tumours are heterogenous displaying several subpopulations with differences in morphology, inflammatory infiltrates, mutational status, and gene expression profiles. In our previous research and that of others, we have shown using immunohistochemistry that in humans and mice, 5hmC is abundant in all the differentiated colonocytes and at very low levels in the colon stem cell compartments. In bulk sequencing therefore, most of the 5hmC comes from the differentiated colonocytes in normal colon, and the global loss of 5hmC in colon tumours seems to occur regardless of the subtype of tumour. Indeed, global 5hmC levels doesn’t stratify between different tumour types. It could be that within a heterogeneous colon tumour a subset of cells, that have retained 5hmC may be more metastatic and be expanded during metastasis and that this clonal outgrowth is what we detect in the liver metastasis.

Building upon our previous hypothesis that the distribution of 5hmC could be an “imprint” of cell identity that is preprogrammed in progenitor cells and established during differentiation [36], we propose that 5hmC does not accumulate in tumour cells, as in progenitor cells, but that the preprogrammed 5hmC imprint is maintained and can be reestablished upon progression to metastasis. Our data showing the similarity between metastatic tumours and normal colon cells would fit this model, with the caveat that malignant cells are epigenetically aberrant. Thus, the model would need to include the additional de novo peaks and take into consideration tumour heterogeneity.

The detection of several loci where 5hmC was recovered at the same site as in the normal colon and then spreading in *cis,* may either be a consequence of additional transcription factors being recruited to these sites, or a lack of factors that normally prevent the spread of 5hmC. The transcription factor binding sites identified at the “recover and spread” regions include SALL4, PAX5, ZNF770 and ZNF121. PAX5 has previously been listed as a transcription factor that binds to unmodified CpG sites and recruits TET-histone modifying complexes [60]. However, it is expressed at very low levels in normal colon and reduced even further in tumours, so unlikely to have a 5hmC memory function in these cells. SALL4 has previously been shown to bind to 5hmC, and to accelerate demethylation through interaction with TET2 [61]. But *SALL4* transcript levels are low in colon and colon tumours, even after being increased in primary tumours. Not much is known about ZNF770 and it has not previously been associated with TET activity or 5hmC levels. In our study *ZNF770* transcript levels were inversely correlated with 5hmC levels in the primary tumour and metastasis. Both ZNF770 and ZNF121 (also known as ZHC32; ZNF20; D19S204) belong to the krüppel C2H2-type zinc-finger protein family. ZNF121 has been shown to be a c-MYC-interacting protein with functional effects on MYC and cell proliferation in breast cancer [62]. C-MYC has also been listed as a transcription factor that preferably binds to its binding sites when the CpGs are unmethylated and recruits TET-histone modifying complexes to maintain the unmethylated state [60]. Both ZNF770 and ZNF121 have TET2 binding sites in their promoters and can be regulated by TET2. Future functional analyses will reveal whether ZNF770 and ZNF121 have binding affinities for 5hmC or 5mC and whether they interact with TET enzymes to regulate 5hmC accumulation in normal tissue or cancers.

Finally, we used a primary colon adenocarcinoma cell line, SW480 as a model system to study 5hmC dynamics in zebrafish migration assays. Despite an ongoing debate regarding the validity of cancer cell migration and invasion from the yolk sac and whether this is an active migration or passive diffusion process [63], zebrafish larvae continue to provide an excellent alternative to mouse xenograft experiments enabling larger throughput, a quicker timeframe, and more replicate experiments. In our assays depleted TET expression significantly reduced the incidence of cell migration which is the first essential epithelial to mesenchymal transition stage of metastasis. The levels of 5hmC in the triple knockout SW480 cells could be rescued by transient ectopic expression of mouse Tet2 constructs. The rescued cells showed increased migration rates in the zebrafish assay.

## Conclusion

The 5hmC profiles from CRC patients together with Kaplan-Meier survival curves, integrative - OMICs analysis and experimental cell biology provide evidence of a biphasic profile of reduced 5hmC in primary cancer that recovers during metastasis. This supports a role for TET enzyme activity in reprogramming primary cancer cells towards mesenchymal migrating cells.

## Methods

### Patient samples

Research using patient tumour samples was conducted under the principles of the World Medical Association Helsinki agreement with ethical approval obtained from the Cambridgeshire Local Research Ethics Committee (LREC references 04/Q0108/ 125 and 06/Q0108/307) as previously reported [21, 64]. From these previously reported cohort of 119 CRC patients, available DNA samples included normal colonic mucosa (n=22), taken from tissue some distance away from the tumour, invasive primary carcinoma (n=65), liver metastatic deposits (n=32) and normal liver (5), taken from a site adjacent to the tumour. For this study 16 patients were selected from the CRC cohort and used for mass spectrometry analyses, and 5 of these were sequenced for 5hmC (Table 1). Under LREC, researchers were not provided with extensive patient data, beyond Dukes stage, age, sex and MSI status where known). Tumour purity for the liver metastsis samples used for 5hmC hMeDIP sequencing, was estimated from DNA sequence data using an AITAC pipeline (REF), which estimates tumour purity based on CNV. AITAC values are given in Table 1.

### Antibodies

Anti-5hmC (RRID:AB_10013602, Active Motif, 39769), anti-TET1 (RRID AB_2537831, Invitrogen, MA5-16312), anti-TET2 (RRID AB_2687506, Sigma-Aldrich, MABE462), anti-TET3 (RRID AB_11150700, GeneTex, GTX121453) and anti-beta tubulin (RRID AB_1841238 Sigma-Aldrich, T8328), anti-mouse IgG (RRID *AB_258476,* Sigma-Aldrich, A2304) and anti-rabbit IgG (RRID AB_10709927, abcam, 205718).

### PCR primer sequences

*TET1*: Fwd 5’CAGATTAGTCAGGAAGGAAGATGTAA3’, Rev 5’ATTTTCCAGGGCTTAAAGTCTTGA3’ ; *TET2*: Fwd 5’GCAGCACACCCTCTCAAGATT3’, Rev 5’AATTCAGCAGCTCAGTCCCTTACT3’ ; *TET3*: Fwd 5’AGAACCAGGTGACCAACGAG3’, Rev 5’CGCAGCGATTGTCTTCCTTG3’; *ZNF121*: Fwd 5’TTCGCCTTTATCGTGGTG 3’, Rev 5’AATGTTGTTGAGGTGCTGAC3’; *ZNF770*: Fwd 5’CCTCAATACCGCCAAGGTCTTTC3’, Rev 5’CCAATGTTGCCTCAAGGCTG3’ ; *SALL4*: Fwd 5’TCGTCTGC TAGCGCTCTTCAGATC3’, Rev 5’CGGCGGGCTGAGTTATTGTTCG3’ *PAX5*: Fwd 5’ GCGCAAGAGAGACGAAGGT3’, Rev 5’CTGCTGCTGTGTGAACAAGTC3’

### Mass spectrometry

1 μg of genomic DNA was incubated with 5 U of DNA Degradase Plus (Zymo Research) at 37°C for 3 h and filtered through Amicon 10 kDa centrifugal filter units (Millipore). The concentrations of 2′- deoxycytidine, 5- methyl-2′-deoxycytidine and 5-hydroxymethyl-2′-deoxycytidine in the filtrate were determined using an AB Sciex Triple Quad 6500 mass spectrometer fitted with an Agilent Infinity 1290 LC system and an Acquity UPLC HSS column. The global levels of 5mC and 5hmC were expressed as percentages over total 2′- deoxycytidines.

### Survival curves

The online SurvExpress tool, Biomarker Viewer (bioinformatics.mx) designed by [65], was used to generate Kaplan Meier curves for time (years) to metastasis. Data from the Loboda Yeatman study of Colon cancer (GSE28722) contained 129 colon cancer patient RNA profiles from a spectrum of clinical stages.

### Immunoblot (Dotblot) for global 5hmC/5mC detection

2 µg of DNA per sample and standards (10 ng cytosine, 5 ng 5-methylcytosine, 0.125 ng 5- hydroxymethylcytosine, Zymo Research) were added to a total volume of 100 µL 0.1M NaOH and denatured at 95°C for 5 min. Then, an equal volume of ice cold 2 M ammonium acetate was added before transfer to a nitrocellulose membrane using a 96-well vacuum dot blot apparatus (GE Healthcare). After crosslinking the DNA onto the membrane using an ultraviolet crosslinker (UVP), total DNA was visualised with methylene blue stain (0.05% methylene blue in 0.3M sodium acetate). The membrane was washed twice in 75% ethanol, blocked in 5% w/v milk in PBST (0.1% v/v Tween-20 in PBS) and incubated with the primary 5hmC antibody (1:5000, Active Motif, 39769) at 4 °C overnight. After three 5 min washes in PBST, the secondary antibody (1:2000, Abcam, ab205718) was applied for detection by chemiluminescence (Thermo Scientific) according to the manufacturer’s instructions. A Fusion SL analyser (Vilber) was used to visualise the signal.

### hMeDIP-seq

Illumina libraries were prepared before the pull-down using 1 to 3 µg of sonicated genomic DNA (Bioruptor). Libraries were prepared using the TruSeq DNA sample preparation kit (Illumina) following manufacturer’s instructions. Adaptor modified genomic DNA was then immunoprecipitated following as decribed by [21]. Input and pull-down material was whole genome amplified as previously described [21].

### Read processing, peak calling and normalisation

hMeDIP-seq reads in FASTQ format were quality checked with FASTQC and trimmed using FASTP in default settings. Processed hMeDIP-seq reads were aligned to the GRCh38.p14 human genome with BOWTIE2 using default settings. Then GreyListChIP was run on input files available to filter out regions with a high signal in the input. Next, MACS2 *callpeak* was then used to call peaks in narrowpeak mode with the following settings: *-g 3.0e9 -B -q 0.01 --bw 300*. Output from MACS2 was normalised with MAnorm2 [17]. Normalised peaks were retained only if they were present in greater than 75 % of the samples in each category.

### Principal Component Analysis

Principal component analysis was performed with the fviz_pca_ind function from the FACTOEXTRA R package.

### 5hmC peak enrichment and Upset plots

MAnorm2 *diffTest* was used to compare two tissue types to each other and find differentially enriched peaks from the MACS2 peaks. Differentially enriched peak counts from MAnorm2 *diffTest* were extracted and plotted using upsetplot.

### Differential expression analysis

RNA-seq FASTQ reads down-loaded from the NCBI Sequence Read Archive (SRA), SRR2089755, deposited by Lee et al. [26]. Patients in this data set ranged from 62-73 in age, were microsatellite stable (MSS), M1, and chemo naïve when sampled, similar to our patient cohort. FASTQ reads were quality checked with FASTQC and trimmed using TRIMMOMATIC. Processed RNA-seq reads were then aligned to the GRCh38.p14 human genome using STAR (v2.7.10). HTseqcount was used to obtain read counts for each gene using the GRCh38.p14 human genome gene annotation. Differential expression was performed with edgeR. Lowly expressed genes were first filtered out using the edgeR function filterByExpr() followed by calcNormFactors() and estimateDisp(). Output from these were then input into a generalised linear model using glmFit().

#### DNA methylation analysis

We downloaded the MBD-seq peaks [27] for colon tumour and liver metastasis tissue and aligned to the GRCh38.p14 human genome. The patient cohort was MSS, CIMP -ve, and had liver metastasis samples from chemo naïve patients. Metastasis samples were not matched to primary CRC donors, were a mix of M0 and M1.

### Extracting 5hmC gain and recover marks

A custom script was created (adjacent_methylation_peaks.py) to extract 5hmC regions that have recovered and spread. This script first filtered out peaks that were absent in over 50% of the samples from the output.narrowPeak files from MACS2 (v2.2.6). It then merged peaks that were within 500 base pairs of each other using bedtools merge. The merged peaks for LT were retained if they were at least 1000 base pairs long, for NC and TC all merged peaks were retained. bedtools intersect was then used to uncover peaks that were present in NC and LT but not TC (recovered) and LT only (gained).

We also performed a similar peak filter for methylation marks to uncover LT 5hmC regions that were near to 5mC in LT. Here, 5mC peak region filtering was also performed similarly to above, using MACS2 output.narrowPeak files. Again a 50% consensus filter was applied for a peak to be retained. Similar to above, LT 5mC peaks were merged if they were within 250 base pairs of each other and retained if they were at least 250 base pairs long. Finally, 5hmC recovered or gained peaks were searched for being with 750 base pairs of a 5mC peak region.

To determine whether the recovery and spreading that we observed was not due to chance we ran a permutation test [66] by randomly shuffling the NC and TC merged peaks within the genome, while leaving the original LT merged peaks. We then calculated the number of recovered and spread peaks (shuffled count), the same as described previously. This was repeated 1,000 times to generate an empirical null distribution of overlap counts and from this we computed p = {number of permutations where the shuffled count >= the observed count (n = 3149)} / {total permutations + 1}

### Motif enrichment

5hmC spread peaks were searched for transcription factor binding motifs using Simple Enrichment Analysis (SEA) with the HOCOMOCO Human v11 Core motif database from the online MEME suite toolbox (v 5.3.3).

### Cell culture

SW480 cells were obtained from the ATCC (SW480 [SW-480] - CCL-228 |. Congenic triple TET knockout SW480 clones and wild type cells were cultured in Dulbecco’s modified eagle medium (DMEM) supplemented with 10% FBS and 1% Penicillin-Streptomycin solution. All cell lines were cultured in a humidified incubator at 37°C and 5% CO_2_ and periodically sent for in-house mycoplasma testing.

#### TET activity assay

Nuclear extracts from SW480 cells and CRISPR knockouts were purified using the ABCAM nuclear extraction kit (AB 113474). 2ug of nuclear extracts were used in an 8 well strip format colorimetric 5mC-hydrolase TET activity kit (Abcam Cat ∼AB156912) according to manufacturer’s instructions and calibrated against TET standards (supplied with the kit, using dilutions in range of 0.05 – 0.5ng/ul)

### CRISPR-CAS9 mediated Triple TET KnockDown

SW480 cells (genotype (*TET1*(+/+);*TET2*(+/+);*TET3*(+/+/+) were sent to Horizon Discovery for CRISPR-CAS9 gene editing. The parental SW480 cells were first targeted for the *TET2* locus (Horizon gRNA1779) and a single clone with an out-of-frame deletion of *TET2*, was subsequently used to target the *TET3* locus (Horizon gRNA1780). A single clone in which all *TET3* alleles were confirmed to have out-of-frame deletions was used to target the *TET1* locus. (Horizon gRNA1778) Two clones were confirmed to have out-of-frame deletions for both *TET1* alleles. Two triple knock-out (TKO) clones, SW480-TKO-217 and SW480-TKO-6B12 were used in this study. TET expression was compared to WT SW480 cells by western blotting and RTqPCR.

### Transfection

To recover the expression of Tet2 in SW480 TET TKO lines, cells cultured to 75% confluency in 12-well plates before being transfected with 200μg of mTet2 plasmid (FH-Tet2-pEF was a gift from Anjana Rao (Addgene plasmid # 41710 ; http://n2t.net/addgene:41710 ; RRID:Addgene_41710), [67]) using Lipofectamine 2000 according to manufacturer’s instructions. Cells were then incubated for 24 hours prior to use in downstream experiments.

### Zebrafish tumour xenograft assays

Tumour cells were labelled in a 5 μM solution of Vybrant-CM-DiI (Invitrogen V-22888), resuspended in DMEM and injected into the perivitelline space (PVS) of 48 days post fertilisation (dpf) *Casper* (roy^-/-^; nacre^-/-^) embryos. Xenografts were incubated at 33°C for 48 hours, followed by counting of metastatic foci under a fluorescent microscope. Prior to injection the embryos were de-chorionated, anaesthetised with 164 mg/L tricaine (Merck A5040), and mounted onto glass slides with 100 mg/ml low-melting point agarose. A micromanipulator and gas injection system was used to inject the cells into the embryos. SW480 cells seeded into 6-well culture plates and grown to 90% confluency, the media was then removed, and the cells were washed in PBS several times before labelling. Cells were then washed three times in PBS before being labelled in with the DILK stain for 15 minutes at 37°C. After labelling, cells were washed for 5 minutes in PBS three times, before being trypsinised and resuspended to a concentration of 2x10^5^ cells/μl. Labelled cells were stored on ice until injection. After injection, zebrafish embryos were released from the agarose, placed into petri dishes containing fresh embryo medium and were incubated at 33°C for 48hours. The xenografts were then fixed in 4% paraformaldehyde, before being visualised using a confocal microscope. The presence/absence of migratory cells and the number of distant foci were recorded for each embryo. The scoring was done by three independent researchers, two of whom were blinded as to which embryos had been injected with control or test cells.

## Declarations

### Ethics approval for human tissue used

Research using patient tumour samples was conducted under the principles of the World Medical Association Helsinki agreement with ethical approval obtained from the Cambridgeshire Local Research Ethics Committee (LREC references 04/Q0108/ 125 and 06/Q0108/307)

### Consent for publication

Not applicable.

### Availability of data and materials

The datasets generated and analysed during the current study are available in the GEO database repository, BioProject accession PRJNA206436 and GEO Series GSE268934. **URL:** GEO Accession viewer (nih.gov)

### Competing Interests

The authors declare that they have no competing interests.

### Funding Information

UKRI: Medical Research Council London GB Grant numbers MR/T000481/1 and MR/P000711/1; Engineering and Physical Research Council Grant number 259868, Biotechnology and Biological Sciences Research Council 2598658. Cancer Research at Bath CR@B network; The University of Cambridge, Cancer Research UK (CRUK SEB-Institute Group Award A ref10182)

### Authors Contributions

SU-L prepared sequencing libraries, analysed data. BM, DOH, YT performed the bioinformatics analyses; FH, DLR, PM, JP, TD-P, JLR and SU-L produced and analysed genomic and molecular biology data; JLR, TD-P, JS, DG and NN designed, performed and analysed zebrafish experiments, AEKI consulted on clinical analysis and coordinated patient sample selection. AM conceived and designed the study and wrote the manuscript together with BM, DOH and FH. All authors have read the manuscript.

## Acknowledgements

Thank you to Dr Anna Caldwell at the mass spectrometry facility at Kings College London.

## Notes

### Competing Interest Statement

The authors have declared no competing interest.

