## Supplementary Figures 1 - 4 for "Colorectal cancer progression to metastasis is associated with dynamic genome-wide biphasic 5-hydroxymethylcytosine accumulation"

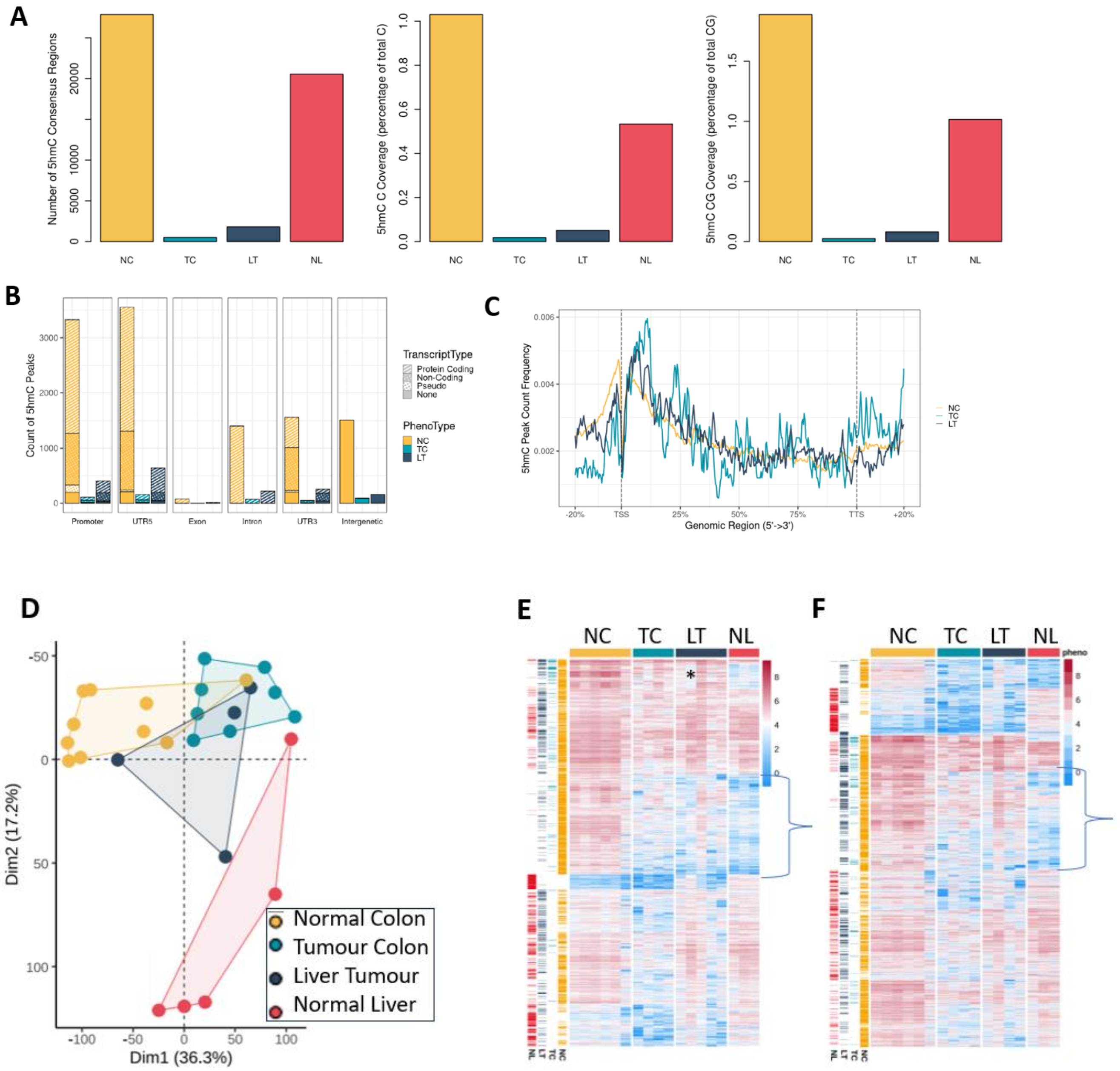


**Figure S1: Total counts of hMeDIPseq peaks (enriched over input) shows metastatic tumours to the liver has more 5hmC enrichment compared to primary tumours, and hMeDIPseq peak loci in metatastic liver tumours cluster towards the colon A)** Number of 5hmC consensus peaks, coverage as a percentage of total C and as a percentage of total CGs. Colour scheme: yellow is normal colon (NC), blue is primary Tumour in Colon (TC), dark blue is metastases to liver tumours (LT) and red is normal liver tissue (NL). **B)** Breakdown of features (Promoters, coding genes, noncoding genes, exons, introns, UTRs at 5hmC peaks **C**) 5hmC count frequency around transcriptional start site (TSS), gene body, and transcriptional termination sites (TSS), for NC (yellow), TC (blue), LT (dark blue). **D-F**) PCA and Heatmap analysis without the potentially contaminated liver tumour sample, mentioned in Fig 2. D) PCA analysis shows tighter clustering towards the colon samples when the outlier sample is removed. E) Heatmap analysis including the contaminated sample (*), F) Heatmap analysis without this sample. The brackets indicate a cluster of loci (Group 4 in Fig 2) within the metastasis sample that have 5hmC peaks in liver tumours similar to normal colon, and which don't have 5hmC in normal liver.

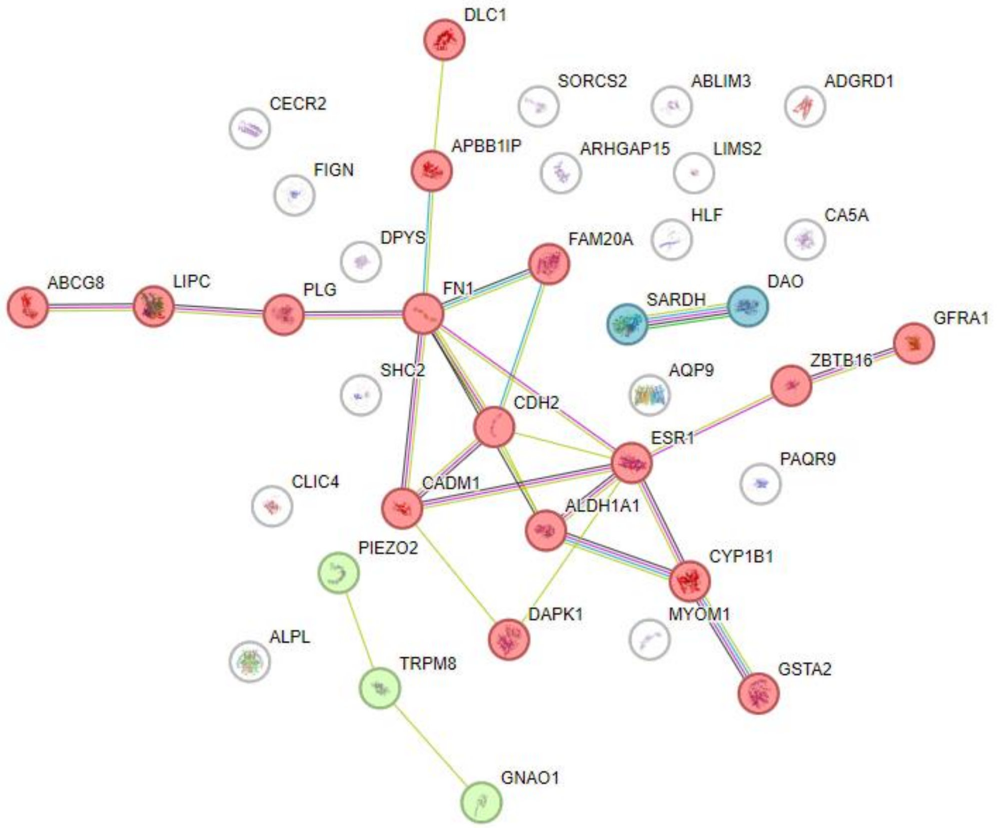


| Biological Process (Gene Ontology) | | | | |
|---|---|---|---|---|
| *GO-term* | *description* | *count in network* | *strength* | *false discovery rate* |
| GO:1904017 | Cellular response to Thyroglobulin triiodothyronine | 2 of 3 | 2.97 | 0.0021 |
| GO:0007161 | Calcium-independent cell-matrix adhesion | 2 of 4 | 2.85 | 0.0028 |
| GO:0033631 | Cell-cell adhesion mediated by integrin | 2 of 6 | 2.67 | 0.0043 |
| GO:1904179 | Positive regulation of adipose tissue development | 2 of 8 | 2.55 | 0.0060 |
| GO:0035357 | Peroxisome proliferator activated receptor signaling pathway | 2 of 9 | 2.5 | 0.0068 |
| GO:0023035 | CD40 signaling pathway | 2 of 10 | 2.45 | 0.0078 |
| GO:0051918 | Negative regulation of fibrinolysis | 2 of 13 | 2.34 | 0.0111 |
| GO:0035313 | Wound healing, spreading of epidermal cells | 2 of 13 | 2.34 | 0.0111 |
| GO:0030949 | Positive regulation of vascular endothelial growth factor rec... | 2 of 14 | 2.3 | 0.0123 |
| GO:0010763 | Positive regulation of fibroblast migration | 2 of 16 | 2.25 | 0.0147 |
| GO:2000811 | Negative regulation of anoikis | 2 of 19 | 2.17 | 0.0194 |
| GO:0033627 | Cell adhesion mediated by integrin | 4 of 42 | 2.13 | 6.88e-05 |
| GO:0035987 | Endodermal cell differentiation | 4 of 44 | 2.11 | 6.88e-05 |
| GO:0034113 | Heterotypic cell-cell adhesion | 3 of 36 | 2.07 | 0.0014 |
| GO:0071634 | Regulation of transforming growth factor beta production | 3 of 39 | 2.03 | 0.0016 |

**Figure S2: STRING analysis of All 5hmC genes with differential expression**. Settings: Interactive sources; High confidence 0.700, 1$^{st}$ shell query proteins only; Kmeans clustering 15. Statistical Background (the whole genome). Full gene list: List: *ABCG8, , ABLIM3, ADGRD1, ALDH1A1, ALPL, APBB1IP, AQP9, ARHGAP15, CA5A, CADM1, CDH2, CECR2, CYP1B1, DAO, DAPK1, DPYS, ESR1, FAM20A, FIGN, FN1, GFRA1, GNAO1, GSTA2, HLF, LINC01348, LIPC, LOC105373215, LOC124904475, MIR99AHG, MYOM1, PAQR9, PIEZO2, PLG, SARDH, SHC2, TRPM8, ZBTB16, LIMS2, DLC1, CLIC4, GNAI, SORCS2, IL6-AS1*, Gene Ontology output with STRING. Table – Gene ontology output of above STRING

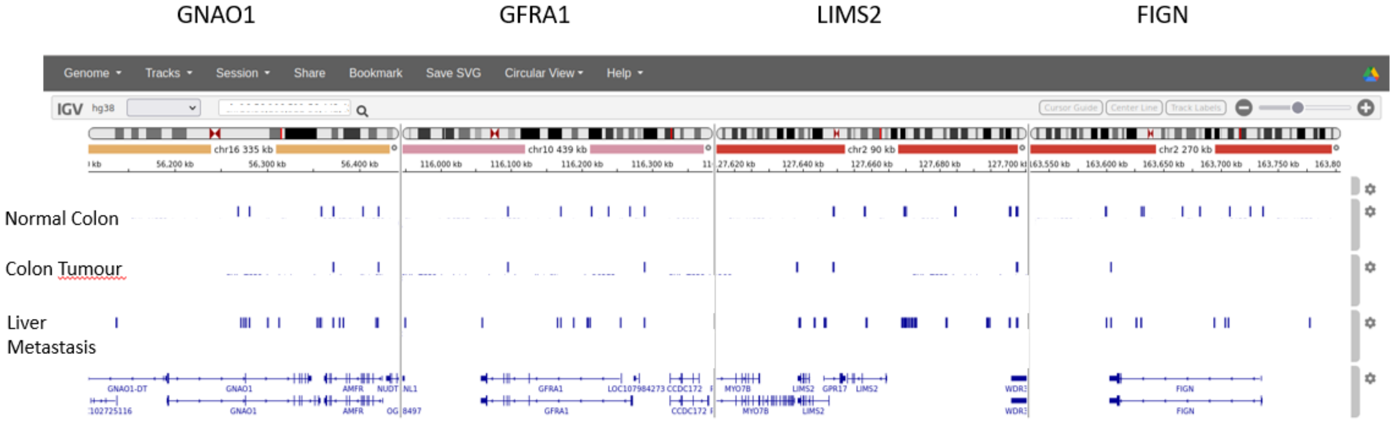


**Figure S3 A visual examination of the 5hmC data on an integrative genomics viewer (IGV)**, showing a pattern of 5hmC "recovering" at sites where 5hmC was present in normal colon tissue, absent in the colon tumour, and recovered in the metastatic tumours.

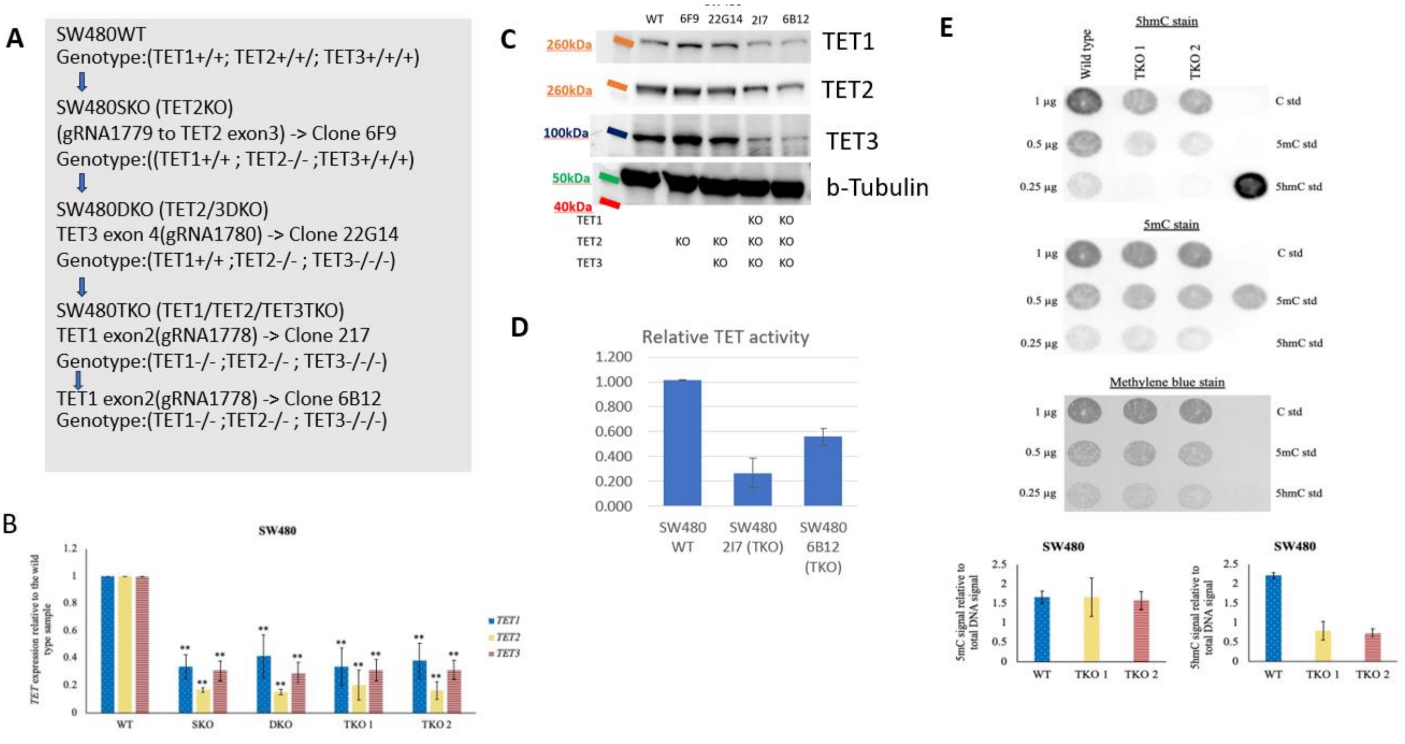


**Figure S4 Effects of Crispr mediated triple TET1/TET2/TET3 knockdown**. A) Schematic of knockdown strategy; B) Effect of knockdown in SW480 cells on TET transcript levels as measured by PCR; C) Western blot analysis to determine TET protein levels after knockdown; D) TET activity in knockout cells relative to wild type measured by colorimetric 5mC hydrolase TET assay kit (ABCAM). E) Immunoblot analyses of 5hmC and 5mC in triple knockout (TKO) and wildtype SW480 cells, with a bar chart below showing quantitative immunoblot confirmation of reduced 5hmC in the two TKO clones.
